# A universal, single primer amplification protocol (R-SPA) to perform whole genome sequencing of segmented dsRNA reoviruses

**DOI:** 10.1101/2021.11.01.466778

**Authors:** Klaudia Chrzastek, Holly S. Sellers, Darrell R. Kapczynski

**Author notes:** Corresponding authors: K.C,; D.R.K.

## Abstract

**Background:** The Reoviridae family represents the largest family of double-stranded RNA (dsRNA) viruses, and the members have been isolated from a wide range of mammals, birds, reptiles, fishes, insects, plants. Orthoreoviruses, one of the 15 recognized genera in the Reoviridae family, can infect humans and nearly all mammals, and birds. Genomic characterization of reoviruses has not been adopted on a large-scale due to the complexity of obtaining sequences for all 10 segments.

**Results:** In this study, we developed a time-efficient, and practical method to enrich reovirus sequencing reads from isolates that allowed for full genome recovery using single-primer amplification method coupled with next generation sequencing. We refer to this protocol as reovirus-Single Primer Amplification (R-SPA). Our results demonstrated that most of the genes were covered with at least 500 reads per base space. Furthermore, R-SPA covered both 5’ and 3’ end of each reovirus genes.

**Conclusion:** A universal and fast amplification protocol that yields double-stranded cDNA in sufficient abundance and facilitates and expedites the whole genome sequencing of reoviruses was presented in this study.

## Background

The flexibility of next-generation sequencing (NGS) technology allows almost any genetic material to be studied on a genome-wide scale. Complete and accurate genome sequencing are essential to facilitate full genome assemblies. Genetic characterization of RNA viruses using sequence-based methods is widely used to understand virus diversity, virus spread, understand origin and evolutionary history of viruses or to perform clinical diagnostics. In eukaryotes, RNA viruses account for the majority of the virome diversity [1]. Orthoreoviruses belong to the Reoviridae family and have a double-stranded RNA (dsRNA) genome. Reoviruses are ubiquitous and can infect humans and animals such as mammals, fish, reptiles and birds. Exposure to reoviruses is presumed to be very common in the human population, 50% of children 5-6 years of age and more than 90% adults can be seropositive to reovirus [2-4]. The orthoreovirus genome consists of ten dsRNA segments that are divided into three classes based on size: large (L1, L2, and L3), medium (M1, M2 and M3) and small (S1, S2, S3 and S4) segments. Each segment encodes one protein, with the exception of segment S1 which contains 3 open reading frames and codes for p10, p17 and σC proteins [5-8]. The segmented nature of the reovirus genome poses a risks for a novel reassortant virus to occur [9-11].

Avian Orthoreoviruses (ARVs) have been isolated from enteric disease syndromes, myocarditis, hepatitis, arthritis/tenosynovitis, malabsorbstion syndrome in commercial poultry and are responsible for economic loss in poultry worldwide [12-14]. Recently, newly emerging avian reovirus variants were isolated in the USA [15-17] or China [18]. The molecular characterization of avian reoviruses is based on amplification of the amino acid sequence of the Sigma C. Sigma C is a major antigenic determinant of ARVs and the most genetically variable gene within the reovirus genome [16, 19-21]. A next-generation sequencing method for avian reoviruses is largely dependent on isolation of the virus and in such case a direct based RNA-Truseq or cDNA HiSeq sequencing results in a high amount of contaminating (non-viral) nucleid acid in the sequences generated as well as, low yields of reovirus-specific sequences upon sequencing. Furthermore, direct sequencing usually requires a multiple template preparation steps such as RNase and/or DNase treatment to remove contaminations, or high-speed sedimentation to concentrate packaged viral genomes. In contrary, targeted reovirus sequencing usually requires the use of multiples reaction to separately target each reovirus gene [18].

In this study, we developed a simple template enrichment protocol that is utilizes one universal primer to target all ten segments of the reovirus genome. We refer to this protocol as reovirus-Single Primer Amplification (R-SPA) and coupled with next-generation sequencing we obtained full genome sequence of reoviruses. Furthermore, we compared a new R-SPA strategy with our previously described sequence-independent single primer amplification (SISPA) that allows for ssRNA viral genome enrichment to assess whether this method could be also applied to dsRNA reoviruses.

## Results

### Whole genome sequencing and genome coverage

A complete, full genome sequence of reoviruses were obtained using the single primer amplification method presented in this study, coupled with next-generation sequencing. We were able to recover 10 segments of each reovirus genome sequence sequenced in this study using R-8N single primer amplification (R-SPA), or a combination of R-8N and R-Rev-8N and obtained a nearly full genome sequence using K-8N SISPA strategy (Table 2). The primary difference between the amplification methods evaluated was observed to be the number of specific reovirus reads and depth of coverage obtained after amplification and sequencing. The lowest number of reads assembled to the host genome and therefore the highest number of reads assembled to the reference reovirus genome was achieved using the R-SPA method (Suppl. Table 1). The percentage of reads assembled to the reovirus reference genome (after filtering against chicken host genome) was 39% (for Ck/2016 isolate), and 51% (for Ck/2017 isolate) when R-SPA was used for genome amplification, followed by 21% (for Ck/2016 isolate) and 33% (for Ck/2017 isolate) using K-8N SISPA strategy and 35% (for Ck/2016 isolate) and 41% (for Ck/2017 isolate) when R-SPA was combined with a helper primer R-Rev-8N and applied for WGS (Suppl. Table 1). Overall, the highest average depth of coverage was achieved for lambda gene segments (λA, λB, λC), followed by µA, and ơC. R-SPA and R-SPA combined with R-reverse-8N primer covered eight or nine out of 10 segments with a depth of coverage above 1000 reads per bp, respectively. K-8N SISPA strategy allowed for six out 10 genes to be covered with at least 1000 reads per bp (Table 2). Mapping of the sequencing reads to the reference genome after R-8N R-SPA was performed at a mean depth of 4380.8 for λA, 3071.4 for λB, 4576 for λC, 4187.15 for µA, 1344.3 for µB, 6566.55 for µNS, 5144.7 for ơC, 874.85 for ơA, 4674.95 for ơB and 918.15 for ơNS. A detailed of the statistics is shown in Table 2. As compared to K-8N SISPA strategy, R-SPA allowed to achieve over 13x higher mean depth of coverage for ơB segment followed by 8.5x higher mean depth of coverage for µNS segment and 2.5x for λB and µB (Figure 3A). The mean depth of coverage for the Sigma C encoding region of S1, was similar after R-SPA and K-8N SISPA strategy and slightly lower when a helper primer was added to the R-SPA reaction. The addition of the Rev-8N primer to R-SPA did not increase the depth of coverage for most of the genome segments sequenced besides sigma NS gene. The sigma NS segment following R-SPA with the helper primer was sequenced at a very high depth of coverage, more than 10 times higher, compared to R-SPA only (10-70x) and over 8 times (8-21x) higher than the K-SISPA strategy (Figure 3A). Overall, the highest depth of coverage was achieved using R-SPA only, followed by R-SPA with a helper primer and K-8N SISPA strategy. The distribution of reads aligned to the reference reovirus genome is shown in Figure 3B. Our results demonstrated that most of the genes were covered with at least 500 reads per base space, especially when R-SPA was used for reovirus genome amplification. Furthermore, R-SPA covered both 5’ and 3’ end of each reovirus genes (Suppl. Figure 1).

**Table 1.**
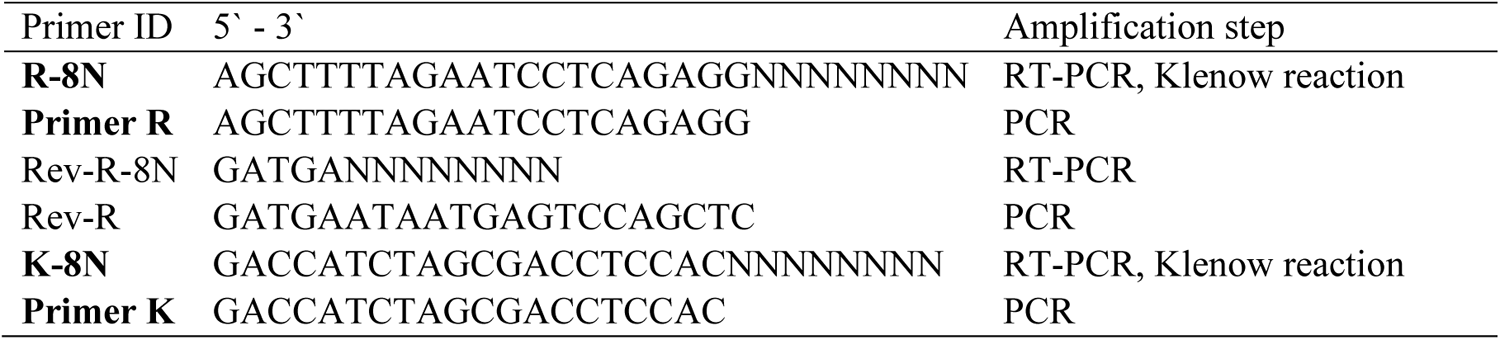
The list of primers used in this study. In bold are the primers that allows for successful whole genome, next-generation sequencing of reoviruses.

**Table 2.**
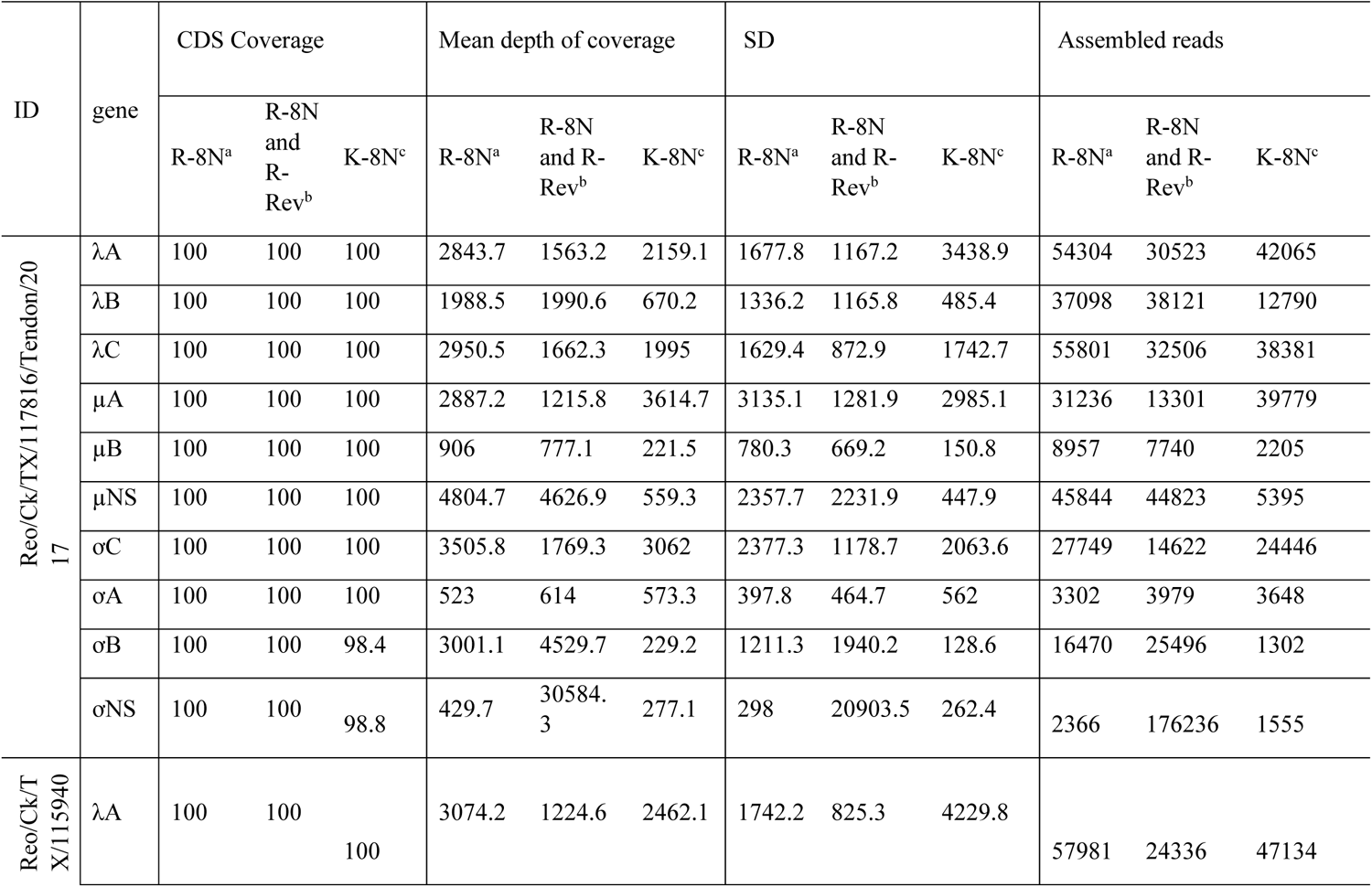
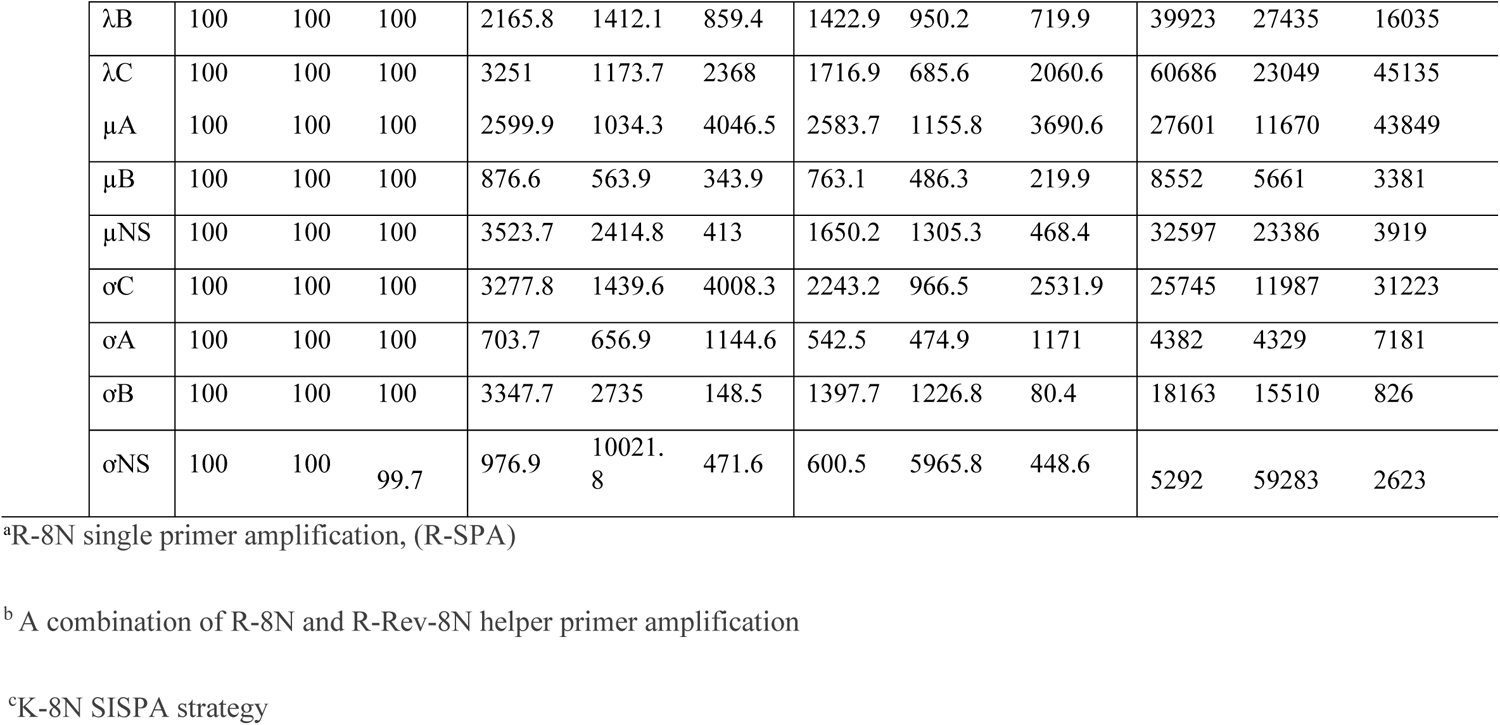
Comparison between amplification methods applied in this study along with Miseq Ilumina sequencing to obtain a full genome of reoviruses.

| ID | gene | CDS Coverage |  |  | Mean depth of coverage |  |  | SD |  |  | Assembled reads |  |  |
| --- | --- | --- | --- | --- | --- | --- | --- | --- | --- | --- | --- | --- | --- |
|  |  | R-8N<sup>a</sup> | R-8N and R-Rev<sup>b</sup> | K-8N<sup>c</sup> | R-8N<sup>a</sup> | R-8N and R-Rev<sup>b</sup> | K-8N<sup>c</sup> | R-8N<sup>a</sup> | R-8N and R-Rev<sup>b</sup> | K-8N<sup>c</sup> | R-8N<sup>a</sup> | R-8N and R-Rev<sup>b</sup> | K-8N<sup>c</sup> |
| Reo/Ck/TX/117816/Tendon/2017 | λA | 100 | 100 | 100 | 2843.7 | 1563.2 | 2159.1 | 1677.8 | 1167.2 | 3438.9 | 54304 | 30523 | 42065 |
|  | λB | 100 | 100 | 100 | 1988.5 | 1990.6 | 670.2 | 1336.2 | 1165.8 | 485.4 | 37098 | 38121 | 12790 |
|  | λC | 100 | 100 | 100 | 2950.5 | 1662.3 | 1995 | 1629.4 | 872.9 | 1742.7 | 55801 | 32506 | 38381 |
|  | μA | 100 | 100 | 100 | 2887.2 | 1215.8 | 3614.7 | 3135.1 | 1281.9 | 2985.1 | 31236 | 13301 | 39779 |
|  | μB | 100 | 100 | 100 | 906 | 777.1 | 221.5 | 780.3 | 669.2 | 150.8 | 8957 | 7740 | 2205 |
|  | μNS | 100 | 100 | 100 | 4804.7 | 4626.9 | 559.3 | 2357.7 | 2231.9 | 447.9 | 45844 | 44823 | 5395 |
|  | σC | 100 | 100 | 100 | 3505.8 | 1769.3 | 3062 | 2377.3 | 1178.7 | 2063.6 | 27749 | 14622 | 24446 |
|  | σA | 100 | 100 | 100 | 523 | 614 | 573.3 | 397.8 | 464.7 | 562 | 3302 | 3979 | 3648 |
|  | σB | 100 | 100 | 98.4 | 3001.1 | 4529.7 | 229.2 | 1211.3 | 1940.2 | 128.6 | 16470 | 25496 | 1302 |
|  | σNS | 100 | 100 | 98.8 | 429.7 | 30584.3 | 277.1 | 298 | 20903.5 | 262.4 | 2366 | 176236 | 1555 |
| Reo/Ck/TX/115940 | λA | 100 | 100 | 100 | 3074.2 | 1224.6 | 2462.1 | 1742.2 | 825.3 | 4229.8 | 57981 | 24336 | 47134 |

|  |  |  |  |  |  |  |  |  |  |  |  |  |
| --- | --- | --- | --- | --- | --- | --- | --- | --- | --- | --- | --- | --- |
| λB | 100 | 100 | 100 | 2165.8 | 1412.1 | 859.4 | 1422.9 | 950.2 | 719.9 | 39923 | 27435 | 16035 |
| λC | 100 | 100 | 100 | 3251 | 1173.7 | 2368 | 1716.9 | 685.6 | 2060.6 | 60686 | 23049 | 45135 |
| μA | 100 | 100 | 100 | 2599.9 | 1034.3 | 4046.5 | 2583.7 | 1155.8 | 3690.6 | 27601 | 11670 | 43849 |
| μB | 100 | 100 | 100 | 876.6 | 563.9 | 343.9 | 763.1 | 486.3 | 219.9 | 8552 | 5661 | 3381 |
| μNS | 100 | 100 | 100 | 3523.7 | 2414.8 | 413 | 1650.2 | 1305.3 | 468.4 | 32597 | 23386 | 3919 |
| σC | 100 | 100 | 100 | 3277.8 | 1439.6 | 4008.3 | 2243.2 | 966.5 | 2531.9 | 25745 | 11987 | 31223 |
| σA | 100 | 100 | 100 | 703.7 | 656.9 | 1144.6 | 542.5 | 474.9 | 1171 | 4382 | 4329 | 7181 |
| σB | 100 | 100 | 100 | 3347.7 | 2735 | 148.5 | 1397.7 | 1226.8 | 80.4 | 18163 | 15510 | 826 |
| σNS | 100 | 100 | 99.7 | 976.9 | 10021.<br>8 | 471.6 | 600.5 | 5965.8 | 448.6 | 5292 | 59283 | 2623 |

<sup>a</sup>R-8N single primer amplification, (R-SPA)
<sup>b</sup> A combination of R-8N and R-Rev-8N helper primer amplification
<sup>c</sup>K-8N SISPA strategy

**Figure 1.**
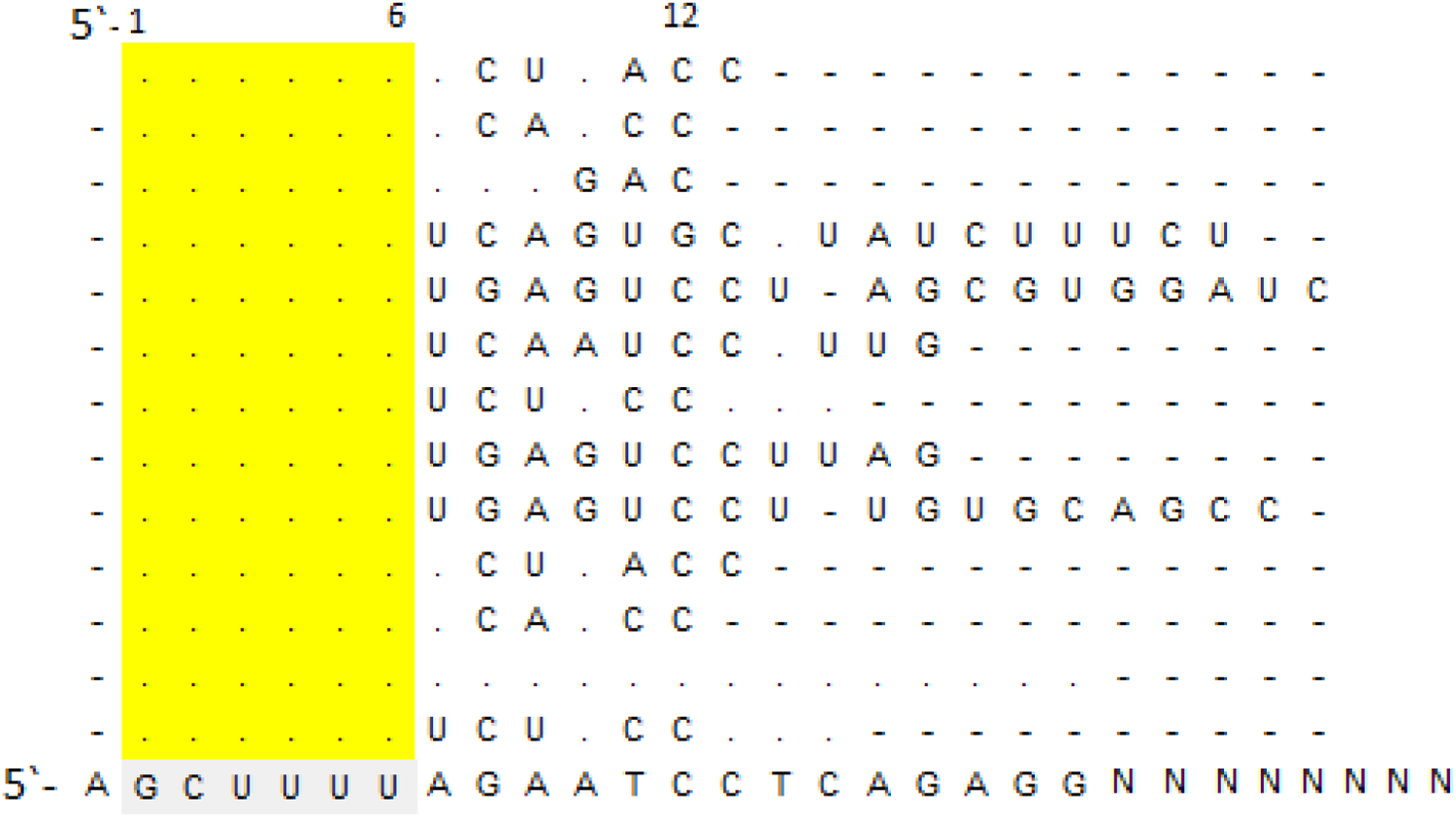
Schematic representation of the conserved 5-end terminal regions of different segments of reoviruses vRNA and the universal primer R-8N designed in this study. For reovirus – Single Primer Amplification (R-SPA), R-8N primer that contains 21 known nucleotides (barcode) tag to random octamer at the 3’ – end was designed. Out of 21 nucleotides, six represent conserved nucleotides found in all reovirus segments (grey color), followed by six nucleotides (5’-end, position 6 - 12) that were commonly found in large viral segments, eight random nucleotides used to increase the annealing temperature and random octamer (8xN).

### Metagenomic detection

Single primer amplification methods presented in this study allowed detection of avian orthoreovirus by metagenomics. For Reo/Ck/TX/117816/Tendon/2017, out of 54,856 reads classified by Kraken, 95% represents avian reovirus (R-SPA strategy), and out of 49,108 reads classified after K-8N SISPA strategy, 77% were assigned to avian reovirus (Table 3). For Reo/Ck/TX/115940/Tendon/2016, 92% and 90% of reads were assigned to avian reovirus after R-8N, R-SPA and K-8N SISPA strategy, respectively (Table 3).

**Table 3.** Metagenomic classification of sequencing reads by Kraken metagenomic classifier (version: 1.0.0).

| Sample ID | Barcode used for amplification | Total reads | Reads classified | The number of reads classified to avian reovirus | The percentage (%) of reads classified to dsRNA viruses |
| --- | --- | --- | --- | --- | --- |
| Reo/Ck/TX/117816/Tendon/2017 | Primer R | 360804 | 54856 | 53294 | 95 |
|  | Primer K | 418857 | 49108 | 31844 | 77 |
|  | Primer R and R-Rev | 338004 | 66047 | 61347 | 96 |
| Reo/Ck/TX/115940/Tendon/2016 | Primer R | 727901 | 88128 | 77993 | 92 |
|  | Primer K | 468743 | 64330 | 53522 | 90 |
|  | Primer R and R-Rev | 352833 | 112487 | 53522 | 90 |

## Discussion

Whole-genome sequencing (WGS) of viral genomes plays a very important role in rapid identification of infection, tracking the source of infection during a disease outbreak, evaluating reassortment events, or tracking the genome changes below the consensus level. To take advantage of WGS, different next-generation methodologies are used for virus sequencing. Sequence-Independent, Single-Primer-Amplification (SISPA) in combination with next-generation sequencing (NGS) was used for characterization RNA and DNA viruses in humans and animals, including influenza virus, human respiratory syncytial virus (HRSV), human metapneumovirus (HMPV), human rhinovirus (HRV) and human parainfluenza virus types 1–3 (HPIV1-3), chikungunya virus (CHIKV), Middle East respiratory syndrome coronavirus (MERS-CoV) [19, 22, 23]. Targeted NGS was implemented for instance for Ebola or chikungunya virus (CHIKV) characterization [24, 25]. We have previously applied SISPA strategy for whole genome sequencing of avian single strain RNA viruses, including avian influenza virus, Coronavirus and Paramyxovirus [26]. In 2012, there were a few reports that regards to a newly emerging reovirus in USA and China in avian species [15, 17, 27]. Furthermore, a novel mammalian orthoreovirus was recently isolated from a child in Japan [28]. Realizing the need for a reliable approach for sequencing reovirus genomes without the need of targeting each segment separately, we have developed and report in this study a methodology that generates complete reovirus genomes by using single primer that is targeting conserved 5′-end of each segment of viral genome followed by Klenow reaction and PCR amplification, which improved the yield of double-stranded cDNA. This product can be then directly used for manual or automated Illumina library preparation workflow. Almost two decades ago, a universal primer set for full-length amplification of all influenza A viruses that similarly like in our study is targeting a conserved 5′- and 3-’ end of segmented influenza genome was developed [29]. Although there was no next-generation sequencing in the beginning of the year 2000, this strategy can serve as a useful indication to design universal primer sets for other segmented viruses as we successfully applied to reoviruses in this study. In this study, we were able to obtain full gene sequences for reoviruses with a high depth of coverage that may be used for a subconsensus-level characterization of viral segments. Furthermore, we have shown that modification of a previously described SISPA NGS strategy could also be successfully applied to amplify the reovirus genome. However, the number of reovirus sequencing reads obtained and depth of coverage after K-8N SISPA strategy was lower than after reovirus-Single Primer Amplification (R-SPA). This indicates that the method of choice should greatly depend on the purpose of the study. For instance, for the purpose of fast clinical diagnostics of an unknown sample, the K-8N SISPA strategy may be useful, whereas the R-SPA strategy would be more beneficial for reovirus characterization.

## Conclusions

In conclusion, a universal and fast amplification protocol that yields double-stranded cDNA in sufficient abundance and facilitates and expedites the whole genome sequencing of reoviruses was presented in this study. Further, our method reduces contamination with non-reovirus sequences that may overwhelm the final sequence output, which can occur using non-selective nucleic acid amplification procedures prior to sequencing. We additionally provide evidence to suggest that the method described is more simplistic than other targeted-based sequencing approaches and could produce ten segments of reovirus genome using only one universal primer. It can be applied in any basic molecular virology/microbiology laboratory with access to a thermal cycling machine.

## Methods RNA isolation

The two reovirus field isolates Reo/Ck/TX/117816/Tendon/2017 (Ck/2017) and Reo/Ck/TX/115940/Tendon/2016 (Ck/2016) used in this study were isolated from chickens. The sequences were submitted to the GenBank under the accession number MT995733-MT995742 (Ck/2017) and MW002447-MW002456 (Ck/2016). The raw reads were deposited under the project number PRJNA662570. Total RNA extraction from virus stock was performed using RNeasy Mini Kit (QIAGEN, Valencia, USA) according to manufacturer’s instruction.

### Design of universal oligonucleotides for reovirus Single Primer Amplification (R-SPA), whole genome sequencing (WGS)

The reovirus genome is composed of ten viral RNA segments. The 5′ and 3′ terminal sequences of reovirus segments were conserved among all sequences aligned in this study. The 5′-end contains six highly conserved molecules (GCUUUU[U/C] whereas 3’-end contains five conserved nucleotides (UCAUC). For reovirus – Single Primer Amplification (R-SPA) we designed “R-8N” primer that contains 21 known nucleotides (barcode) tag to random octamer at the 3’ – end. Out of 21 nucleotides, six represent conserved nucleotides found in all reovirus segments, followed by six nucleotides (5’-end, position 6 - 12) that were commonly found in large viral segments, and eight random nucleotides used to increase the annealing temperature (Figure 1). A “Reverse R-8N” (R-Rev-8N) and “Reverse-R” (Rev-R) primers were designed as helper primers to target 3’-ends of template in R-SPA strategy. The list of primers used in this study is shown in Table 1.

### RT-PCR

Schema of sample amplification is shown in Figure 2. For cDNA synthesis a new primer, R-8N, was designed and compared with the previously described primer K-8N [26] and the combination of primers R-8N and helper primer R-Rev-8N using single primer amplification strategy. Extracted RNA (5 μl) was heat treated at 95°C for 4 minutes in the presence of 1 μl of 100 μM primer R-8N (Method 1), 1 μl of 100 μM of K-8N (Method 2), or 1 μl of 100 μM primer R-8N and 1 μl of 100 μM primer R-Rev-8N (Method 3) and dNTPs (10 μM each) in 20-μl reaction mixture. Reverse transcription with SuperScript IV Reverse Transcriptase (ThermoFisher Scientific) was performed at 55°C for 10 min, followed by 80°C for 10 min and then placed on ice.

**Figure 2.**
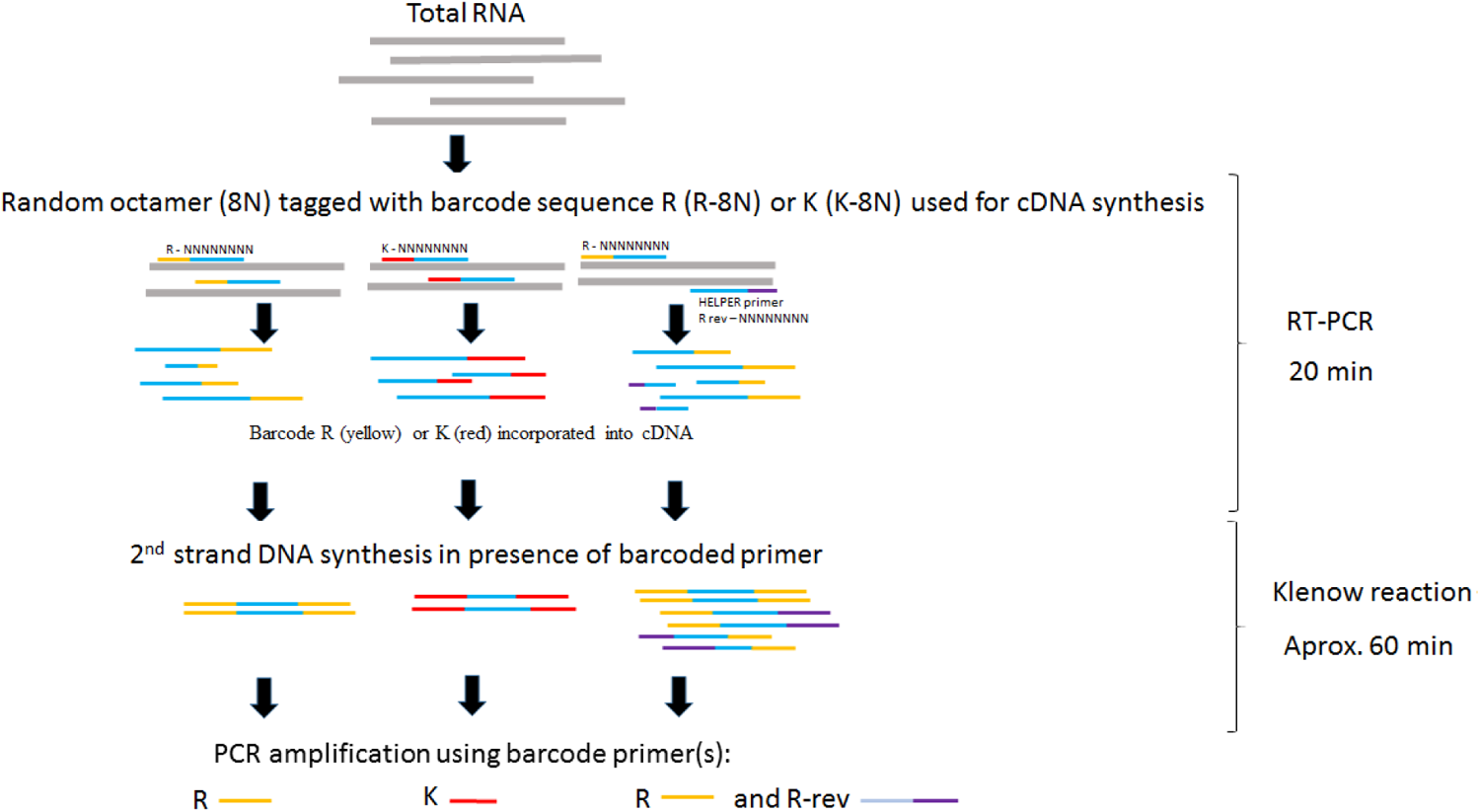
A schema of reovirus amplification. Total RNA was extracted using commercially available kit. A reovirus - Single Primer Amplification, R-SPA protocol consists of three main steps: (i) RT-PCR reaction, (ii) Klenow reaction and (iii) conventional PCR reaction. In the first step, single primer (barcode) that is tagged to random octamer is used to convert RNA into cDNA (R-8N, yellow; K-8N, red). Next, cDNA is converted into dsDNA by Klenow polymerase in a presence of K-8N or R-8N primer in an isothermal amplification. The purified dsDNA of the Klenow reaction is subsequently used as a template for PCR amplification using barcode primer (primer R or primer K). After PCR amplification, and size selection using Agencourt AMPure XP beads (Beckman Coulter) the product can be used as an input for libraries preparation and sequencing.

**Figure 3.**
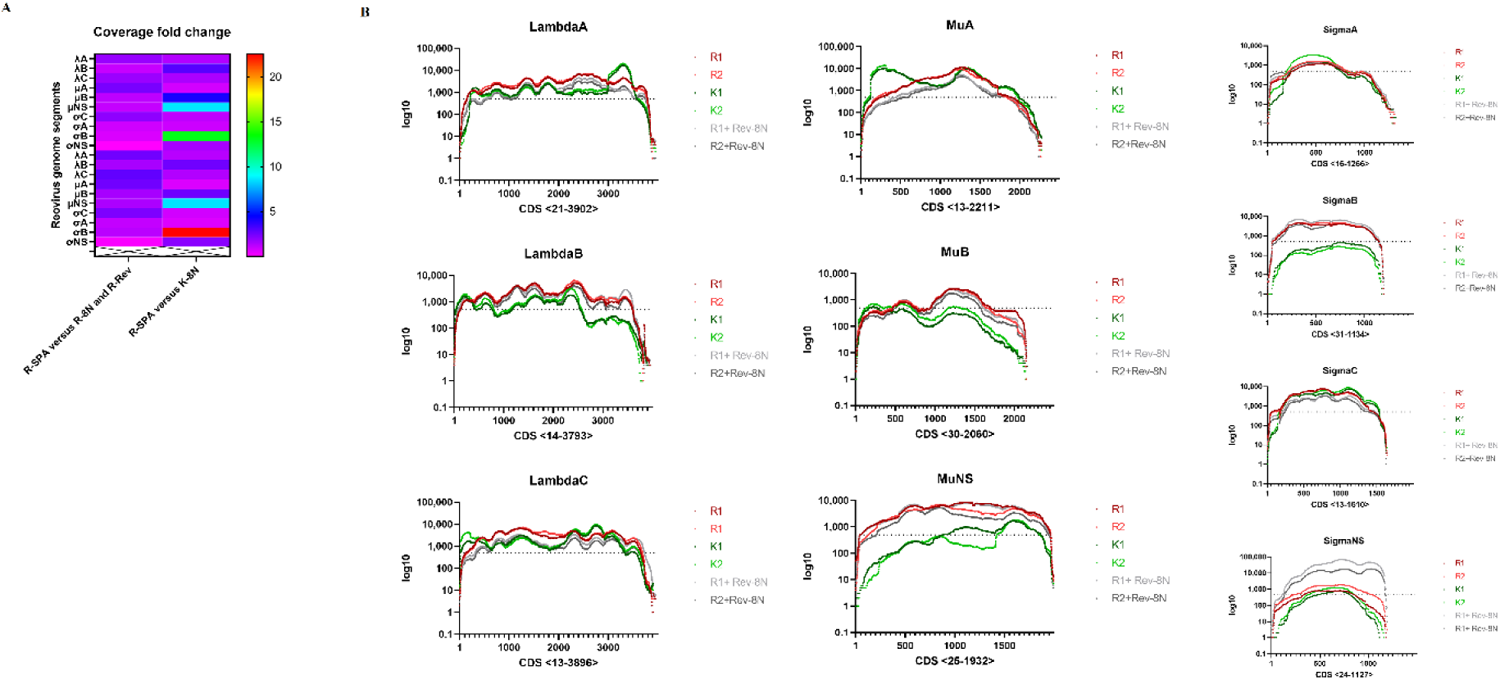
A. Coverage fold changes of reovirus genome after R-Single Primer Amplification (R-SPA) coupled with next-generation sequencing as compare to R-SPA with R-Rev primer amplification and K-8N SISPA strategy. B. Distribution of aligned reads across the reovirus genome. Each graph represents distribution of reads achieved after R-Single Primer Amplification (R-SPA) (green, R1 and violet, R2), K-8N SISPA strategy (K1, black and K2, red) and R-SPA in combination with helper primer R-Rev (grey, R+Rev1 and salmon, R+Rev 2). The CDS position of each gene is indicated below the graph. Grey line represents at least 500 of reads aligned.

To convert the first-strand cDNA into double-stranded cDNA, 20 μl of the first-strand cDNA was heated to 94 °C for 3 min and then cooled to 4 °C in the presence of 0.5 μl of 10 μM either primer R-8N, or 0.5 μl of 10 μM of K-8N, or combination of primer R-8N and 0.5 μl of 10 μM primer R-Rev-8N and dNTPs (10 μM each) in 1× Klenow reaction buffer (NEB). Afterwards, 1 ul of Klenow fragment (NEB) was added and incubated at 37 °C for 60 min (final volume, 25 μl). After conversion into dsDNA by Klenow polymerase (NEB), the products were purified using Agencourt AMPure XP beads (Beckman Coulter). The purified dsDNA of the Klenow reaction was subsequently used as a template for PCR amplification.

### PCR amplification

PCR amplification was conducted with 5 μl of the double-stranded cDNA template in a final reaction volume of 50 μl, which contained 1× Phusion HF buffer, 200 μM deoxynucleoside triphosphate (dNTP), 0.5 U Phusion DNA polymerase (NEB) and 2.5 μl of 10 μM either primer R (Method 1), or 2.5 μl of 10 μM primer K (Method 2) or 2.5 μl of 10 μM primer R and 2.0 μl of primer R-Rev (Method 3). The PCR cycling was performed as follows: 98°C for 30 s, followed by 35 cycles of 98°C for 30 s, 50°C for 30 s, and 72°C for 1 min, with a final extension at 72°C for 10 min. PCR products were purified using Agencourt AMPure XP beads (Beckman Coulter) (in ratio 0.7x). For quantification of the ds cDNA, the Qubit dsDNA HS assay (Invitrogen) was performed according to the manufacturer’s instruction.

### Genome sequencing

A total 1 ng of DNA was used to prepare the library using the Nextera XT Fragment Library Kit (Illumina). Libraries were analysed on a High Sensitivity DNA Chip on the Bioanalyzer (Agilent Technologies) before loading on the flow cell of the 500 cycle MiSeq Reagent Kit v2 (Illumina, USA) and pair-end sequencing (2 × 250 bp). The whole-genome sequencing was performed using the Illumina MiSeq platform.

### Data analysis

The quality of sequencing reads was assessed using FastQC ver. 0.11.5. The reads were then quality trimmed with Phred using a quality score of 30 or more, in addition to low-quality ends trimming and adapter removal using Trim Galore ver.0.5.0 (powered by Cutadapt) ((https://github.com/FelixKrueger/TrimGalore). Filtering the reads against host genome (Gallus gallus 4.0) was performed using BWA-MEM [30]. Unmapped reads from the BAM file were extracted using Samtools 1.9 [31]. The direct assembly of sequencing reads to reference reovirus genome were performed using the Geneious 9.1.2 [32]. The reads were directly mapped to a reference genome Avian orthoreovirus strain Reo/PA/Layer/27614/13 (10 segments, GenBank accession no. KU169288–KU169297). In parallel with reference assembly, Kraken Metagenomics Classifier Version: 1.0.0 [33] was used to analyse sequence sets.

## Declarations

### Ethics approval and consent to participate

Not applicable.

### Consent for publication

Not applicable.

### Availability of data and material

All data generated or analysed during this study are included in this article. The sequences were submitted to the GenBank under the accession number MT995733-MT995742 and MW002447-MW002456. The raw reads were deposited under the project number PRJNA662570.

## Authors’ contributions

K.C, H.S.S, D.R.K contributed to the study conception and design. Material preparation and sequencing was performed K.C. Data curation, KC. Data analysis was performed by K.C, H.S.S, D.R.K. The first draft of the manuscript was written by K.C and all authors commented on previous versions of the manuscript. All authors read and approved the final manuscript.

## Acknowledgements

Not applicable.

---

**Supplementary Table 1.** Basic statistics achieved after single primer amplification of reovirus genome using different primers(barcodes).

| Sample ID | Barcode | Total reads number | Number of reads BWA-MEM mapped to chicken genome (Gallus Gallus 4.0) | Mapped to reference reovirus out of filtered against host genome |
| --- | --- | --- | --- | --- |
| Reo/Ck/TX/117816/Tendon/2017 | R | 725763 | 426727 (58.8%) | 283127 (84%) |
|  | K | 791660 | 612570 (77.38%) | 171566 (92.2%) |
|  | R and R-Rev | 1082164 | 660437 (61.03%) | 387347 (91.85%) |
| Reo/Ck/TX/115940/Tendon/2016 | R | 549521 | 251371 (45.74%) | 280922 (85.64%) |
|  | K | 603870 | 393944 (65.24%) | 201306 (93.11%) |
|  | R and R-Rev | 501915 | 269071 (53.61%) | 206646 (84.72%) |

**suppl Figure 1.**
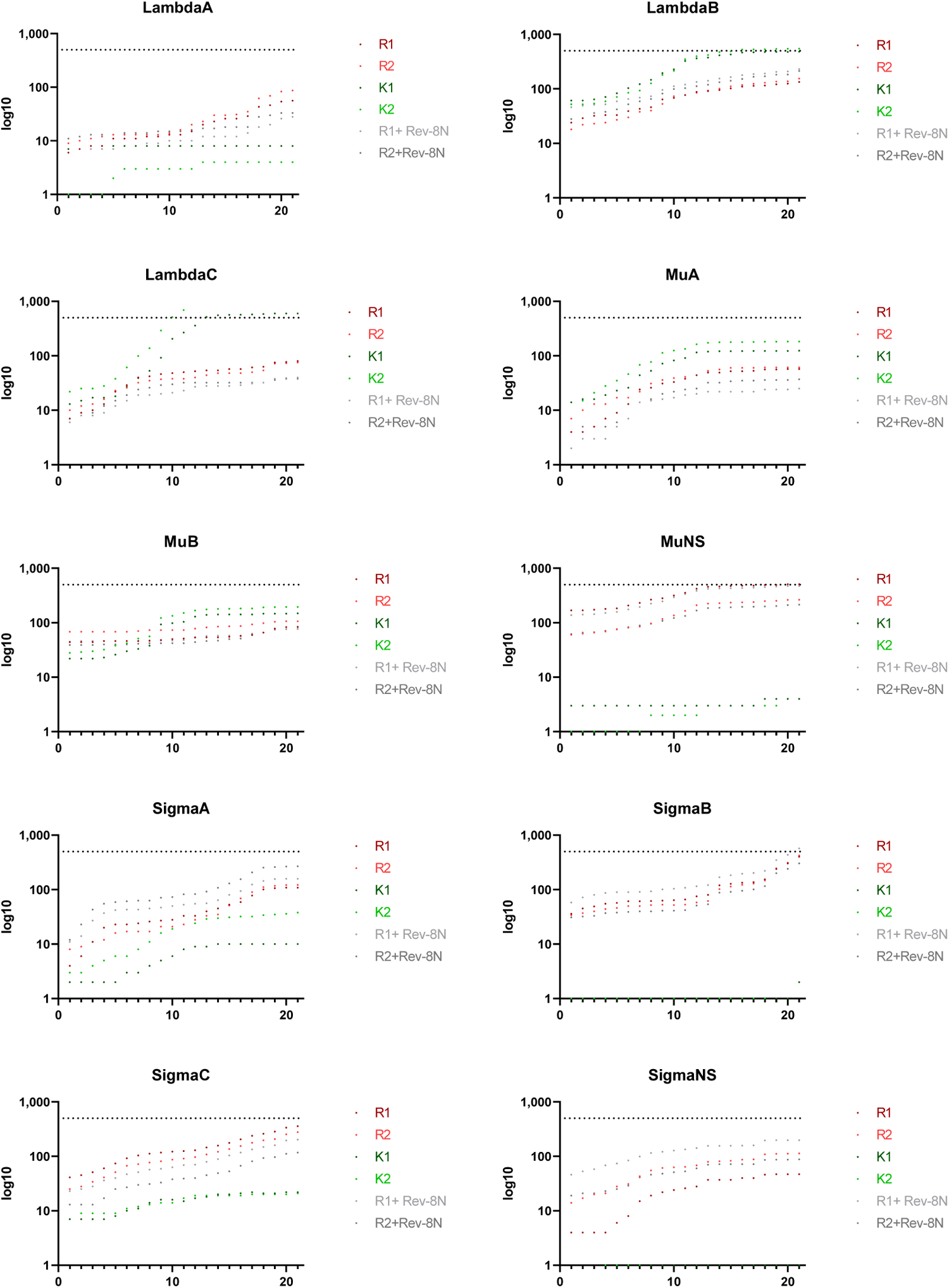
Distribution of aligned reovirus reads at 5’-end. Each graph represents distribution achieved after R-Single Primer Amplification (R-SPA) (red, R1 and salmon, R2), K-8N SISPA strategy (K1 and K2, green) and R-SPA in combination with helper primer R-Rev (grey, R+Rev1 and black, R+Rev 2).

